# CRISPR/Cas9-based Editing of a Sensitive Transcriptional Regulatory Element to Achieve Cell Type-Specific Knockdown of the NEMO Scaffold Protein

**DOI:** 10.1101/450320

**Authors:** Milad Babaei, Yuekun Liu, Shelly M. Wuerzberger-Davis, Alan T. Yeo, Larisa Kagermazova, Ethan Z. McCaslin, Shigeki Miyamoto, Thomas D. Gilmorea

**Author notes:** These two authors contributed equally (Milad Babaei, Yuekun Liu).

## Abstract

The use of alternative promoters for the cell type-specific expression of a given mRNA/protein is a common cell strategy. NEMO is a scaffold protein required for canonical NF-κB signaling. Transcription of the *NEMO* gene is primarily controlled by two promoters: one (promoter B) drives *NEMO* transcription in most cell types and the second (promoter A) is largely responsible for *NEMO* transcription in liver cells. Herein, we have used a CRISPR/Cas9-based approach to disrupt a core sequence element of promoter B, and this genetic editing essentially eliminates expression of NEMO mRNA and protein in 293T human kidney cells. By cell subcloning, we have isolated targeted 293T cell lines that express no detectable NEMO protein, have defined genomic alterations at promoter B, and do not support canonical NF-κB signaling in response to treatment with tumor necrosis factor (TNF). Nevertheless, non-canonical NF-κB signaling is intact in these NEMO-deficient cells. Expression of ectopic NEMO in the edited cells restores downstream NF-κB signaling in response to TNF. Targeting of the promoter B element does not substantially reduce NEMO expression (from promoter A) in the human SNU-423 liver cancer cell line. We have also used homology directed repair (HDR) to fix the promoter B element in a 293T cell clone. Overall, we have created a strategy for selectively eliminating cell type-specific expression from an alternative promoter and have generated 293T cell lines with a functional knockout of NEMO. The implications of these findings for further studies and for therapeutic approaches to target canonical NF-κB signaling are discussed.

## INTRODUCTION

Much of the gene diversity in humans is generated by the use of alternative splicing and alternative promoters (Ayoubi & Van De Ven, 1996: Davulri et al., 2008). It is estimated that over 50% of human genes have alternative splicing and/or use alternative promoters, and alternative promoter usage has also been coupled to alternative splicing (Modrek & Lee, 2002; Davuluri et al., 2008; Xin et al. 2008). In many cases, alternative promoters are used for the tissue-specific or developmentally timed expression of a given gene, and abnormal alternative splicing or promoter usage has been associated with human disease, especially cancer (Davuluri et al. 2008; David & Manley, 2010; Zhang & Manley, 2013; Vacik & Rasda, 2017). For some genes, alternative promoters direct the expression of an identical protein coding region in different cell types or under different conditions by virtue of the promoters being located upstream of distinct 5’ non-translated exons that splice to a common set of downstream coding exons. Methods for assessing the function of tissue-specific alternative promoter usage for individual genes are limited. In this paper, we have used a CRISPR/Cas9-based targeting approach to investigate cell type-specific promoter expression of a key gene (*NEMO*) in NF-κB signaling.

The mammalian NF-κB transcription factor is involved in the regulation of many cell and organismal processes (Hayden & Ghosh, 2012). NF-κB itself is tightly regulated by subcellular localization: that is, NF-κB is located in the cytoplasm when inactive, and is induced to translocate to the nucleus when activated by upstream signals. In canonical NF-κB signaling, NF-κB is activated by IKKβ-mediated phosphorylation of the NF-κB inhibitor IκB, which is then degraded to allow NF-κB to enter the nucleus. In non-canonical signaling, cytoplasmic NF-κB p100 is phosphorylated by IKKα, an event that induces proteasome-mediated processing of p100 to p52, which then enters the nucleus to affect gene expression (Sun, 2011).

NEMO (NF-κB Essential MOdulator) is a protein that serves as a scaffold for IKKβ in canonical NF-κB signaling (Maubach et al., 2017). In the absence of NEMO, canonical NF-κB signaling cannot be activated (Schmidt-Supprian et al., 2000). In contrast, activation of non-canonical processing of NF-κB p100 generally does not require NEMO (Sun, 2011). As such, NEMO is a key regulator for activation of the canonical NF-κB pathway by a variety of upstream signals, and NEMO serves to distinguish activation of canonical and non-canonical NF-κB pathways.

NEMO is also involved in human disease in two prominent ways. First, mutations in the *NEMO* gene (*IKBKG*, chromosome X), which often compromise the ability of NEMO to support activation of NF-κB, lead to a variety of developmental and immunodeficiency diseases in humans (Courtois & Gilmore, 2006; Maubach et al., 2017). Second, NEMO is required for the constitutive and chronic activation of canonical NF-κB signaling that occurs in a variety of cancers and is required for the ability of these cancer cells to proliferate or survive (i.e., avoid apoptosis) (Maier et al., 2013; Maubach et al., 2017; Puar et al., 2018). Therefore, inhibition of canonical NF-κB signaling can inhibit proliferation or induce apoptosis in a variety of cell- and animal-based cancer models (Puar et al., 2018). However, enthusiasm for NF-κB-directed inhibition for cancer therapy was greatly dampened by the finding that systemic and genetic inhibition of canonical NF-κB in animal models leads to liver toxicity and often cancer (Grivennikov et al., 2010; Luedde & Schwabe, 2011). For example, mice with liver-specific knockouts of *NEMO* develop liver damage and sometimes cancer (Luedde et al., 2007; Beraza et al., 2009).

We had three goals in this research: 1) to demonstrate that CRISPR-based targeting of an alternative promoter can be used to knock down expression of a gene in a cell type-specific manner; 2) to create a NEMO-deficient, highly transfectable human cell line for NEMO mutant analysis; and 3) to establish a proof-of-principle concept for targeting the NF-κB signaling pathway for disease therapy in a way that might circumvent unwanted side effects in the liver.

## RESULTS

### CRISPR-based targeting of a core promoter sequence in Exon 1B of the *NEMO* gene abolishes NEMO protein expression in HEK 293T cells

The humanIKBKG gene (*NEMO*, herein), encoding the NEMO protein, has four alternative 5’ non-coding exons (1D, 1A, 1B, 1C) that direct transcription in a tissue-specific fashion (Fusco et al., 2006) (Fig. 1A). Exon 1B is the most commonly used first exon in most cell types, and this region has a strong RNApolII, H3K4-me3 and DNAse hypersensitivity peaks in human HEK 293 cells (Fusco et al., 2006) (Fig. 1A). Moreover, the exon 1B-containing mRNA is part of the major *NEMO* transcript found on polysomes in human 293T embryonic kidney cells (Floor & Doudna, 2016d). Within exon 1B, we noted a sequence (ACCGCGAAACT) that is just downstream of a major transcription start site (TSS) of the *NEMO* gene and that is within a consensus sequence which is located near the TSS of many genes (Vo Ngoc et al., 2017a) (Fig. 1A). Based on these cumulative observations, we put forth the hypothesis that this sequence is important for efficient transcription of the *NEMO* gene in 293T cells.

**FIG 1.**
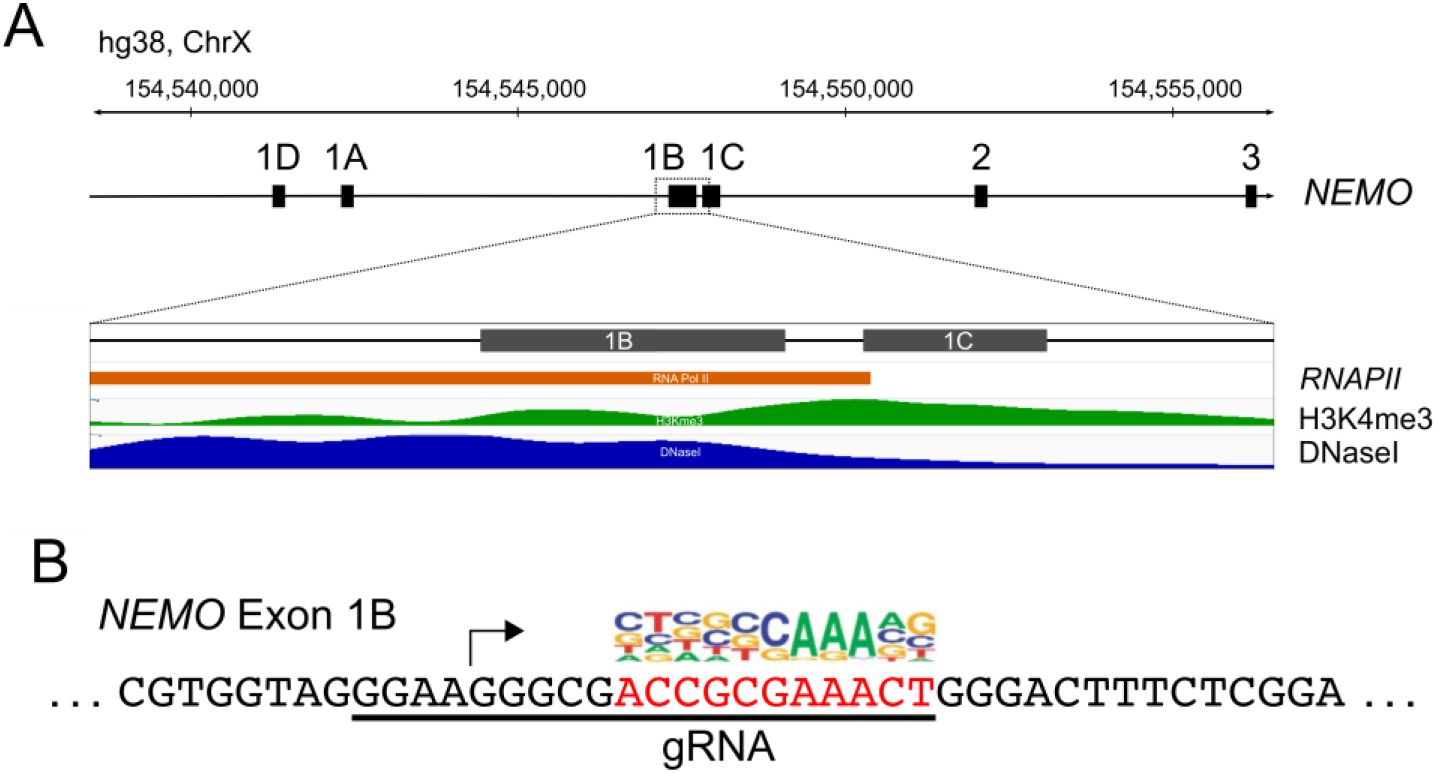
General structure of the 5’ portion of the human *NEMO* gene. (A) Shown are the four 5’ alternative non-coding exons (1D, 1A, 1B, 1C) of the *NEMO* gene on chromosome X, as determined by Fusco et al. (2006). *NEMO* exon 1B has RNAPII, H3K4me3 and DNase hypersensitive site footprints in HEK 293 cells (https://www.encodeproject.org/experiments/ENCSR000DTU/; https://www.encodeproject.org/experiments/ENCSR000EJR/). (B) Downstream of the *NEMO* exon 1B transcription start site (arrow) is a sequence (red) that aligns with a consensus motif (above the red box) that is found near transcription start site of many genes (Vo Ngoc et al. 2017a).

As a first step in testing that hypothesis, we sought to disrupt the predicted exon 1B core promoter element by CRISPR/Cas9 targeting in 293T cells using lentiviral transduction of Cas9 and a gRNA targeting the identified site. After puromycin selection, we performed anti-NEMO Western blotting on extracts from a pool of transduced 293T cells. As shown in Fig. 2A, the levels of NEMO protein were reduced in two independent pools of cells transduced with the lentivirus containing the targeting gRNA as compared to cells transduced with the same vector containing no gRNA. Equal levels of total protein (as judged by β-tubulin Western blotting) were present in both cell lysates, and the FLAG-tagged Cas9 protein was expressed in all transduced cells (Fig. 2A).

**FIG 2.**
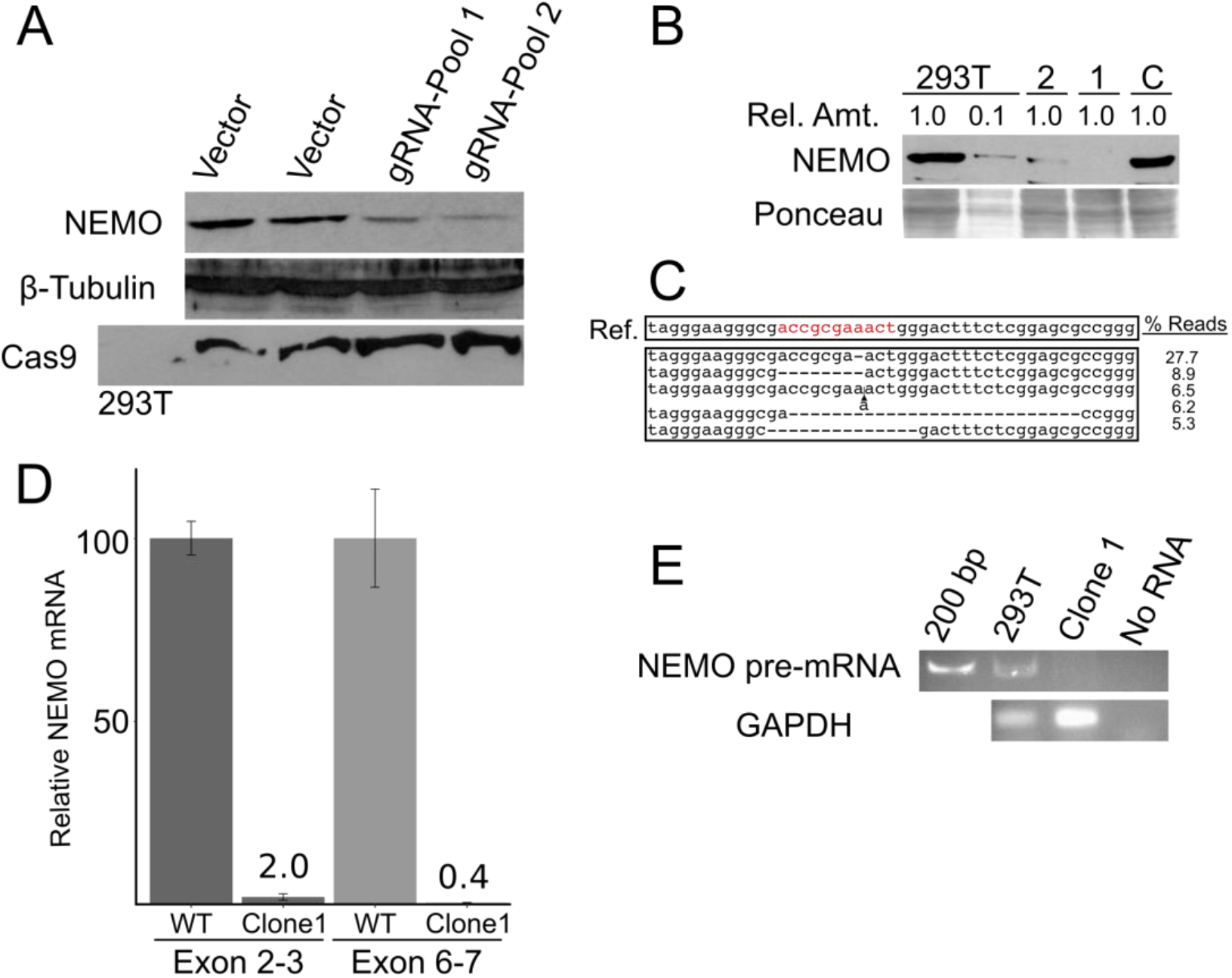
CRISPR/Cas9-based targeting of the exon 1b core promoter disrupts NEMO protein expression in 293T cells. (A) 293T cells that were infected with LentiCRISPR2.0 construct containing no gRNA or the exon 1B gRNA (gRNA-Pool 1 and −2) were selected with puromycin, and puromycin-resistant pools of cells were then subjected to Western blotting for NEMO, β-tubulin (as a loading control), or FLAG-Cas9. (B) Shown is an anti-NEMO Western blot of control 293T cells, a 1:10 dilution of the control extract, and two clones (1 and 2) derived from 293T cells transduced with the LentiCRISPR-exon 1B virus. A Ponceau stain of the filter (as a loading control) is shown at the bottom. As a further control, a clone of puromycin-resistant cells targeted with a different gRNA were analyzed (C). (C) Sequencing of the targeted genomic locus in clone 1 cells was done by PCR amplification of the targeted site and Illumina-sequencing of the PCR product. Shown are the wild-type reference sequence and the five most abundant genomic sequences. The complete array of deletions is shown in Fig. S1. (D) qPCR of *NEMO* transcripts (using exon 2 and 3 or exon 6 and 7 primers) was performed with RNA from control cells and clone 1 cells. The amount of *NEMO* mRNA is relative to the amount of RNA in control 293T cells (100). (E) RT-PCR using primers in exon 1B and the flanking intron 1 was performed with RNA from control and clone 1 cells. Products were then analyzed by gel electrophoresis and staining with ethidium bromide. As a control, shown also is an RT-PCR reaction for *GAPDH* mRNA.

In an effort to identify a clone of cells with a total disruption of the targeted exon 1B promoter sequence, we picked several clones of puromycin-selected cells transduced with the NEMO-targeting lentivirus. Screening of those cell clones by anti-NEMO Western blotting enabled us to identify a cell clone (clone 1) that expressed less than 5% of the NEMO protein expressed in the parental, wild-type 293T cells (Fig. 2B). A second targeted cell clone (clone 2) had clearly reduced levels of NEMO protein, whereas a clone of 293T cells targeted with a gRNA located downstream of exon 1B (clone C, Fig. 2B) did not have reduced NEMO protein expression as compared to parental 293T cells.

To characterize the genomic disruptions in clone 1, we used PCR to amplify the region surrounding the gRNA-targeted site and subjected the pooled PCR product to next generation sequencing. We obtained approximately 34,000 sequence reads from the targeted region in clone 1 cells. All relevant sequences from clone 1 cells had disruptions within the predicted gRNA site, and all had disruptions within the consensus core promoter sequence (ACCGCGAAACT). With a cutoff of 0.26% of total reads, we identified 42 genomic disruptions at the targeted site in clone 1 cells (Fig. S1). The major genomic disruption (comprising 27.7% of the reads) had a one base pair deletion in the non-variant AAA sequence in the consensus sequence (i.e., from ACCGCG<u>AAA</u>CT to ACCGCG_AACT) (Fig. 2C). These results make three points: 1) based on the depth of sequencing, there are few, if any, wild-type exon 1B sequences within clone 1; 2) there is genomic heterogeneity at the targeted site within clone 1; and 3) the lack of NEMO protein in clone 1 is likely due to disruption of the targeted sequence, even by as little as a 1-bp deletion.

We next compared the expression of *NEMO* mRNA in wild-type 293T cells and clone 1 cells. First, we used qPCR to compare the total *NEMO* mRNA in the wild-type and clone 1 cells, by using primer sets downstream of the targeted upstream region, i.e., in exons 2 and 3 or in exons 6 and 7. As shown in Fig. 2D, the total *NEMO* mRNA was reduced by at least 50-fold in clone 1 cells. Similarly, using primers in exon 1B and in the proximal first intron to detect unspliced *NEMO* pre-mRNA, we detected an appropriately sized band in RNA from wild-type 293T cells, but no amplified product when using RNA from clone 1 cells (Fig. 2E). The levels of control *GAPDH* mRNA were similar in both cell types (Fig. 2E). Thus, the levels of pre- and mature *NEMO* mRNA are greatly reduced in clone 1 cells, which have a variety of genomic deletions in a consensus exon 1B core promoter sequence, and these disruptions likely account for the lack of *NEMO* mRNA and consequently NEMO protein expression in clone 1 cells.

### Clone 1 cells are defective for induced activation of NF-κB signaling

To determine whether the lack of NEMO expression renders clone 1 cells defective for NF-κB signaling, we compared the activation of NF-κB signaling in wild-type and clone 1 cells in response to a variety of agents (TNFα, and DNA-damaging agents camptothecin, VP16, doxorubicin, and gamma irradiation). As shown in Fig. 3A, clone 1 cells did not show increased nuclear NF-κB DNA-binding activity in response to any of these agents, whereas control 293T cells showed robust induction of nuclear NF-κB DNA-binding activity. Moreover, induced phosphorylation of IκBα was not detected in clone 1 cells in response to any of these agents (Fig. 3A), but was seen in control 293T cells. As controls, we show that β-tubulin expression and Oct DNA-binding activity are similar in wild-type and clone 1 cells, under all conditions (Fig. 3A).

**FIG 3.**
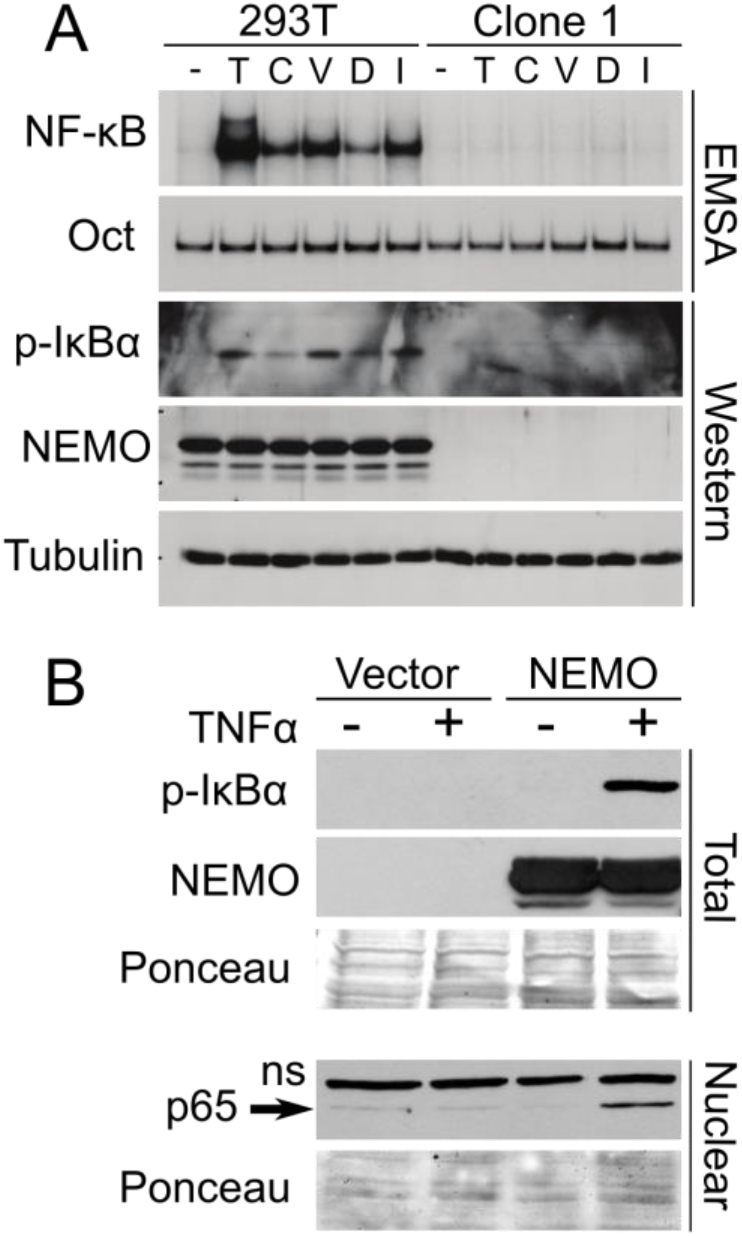
Clone 1 cells are defective for canonical NF-κB activation in response to multiple activators. (A) Control 293T and clone 1 cells were treated with the indicated inducers of canonical NF-κB signaling, and extracts were then analyzed by EMSA for NF-κB and Oct1 DNA binding, or by Western blotting for NEMO, phospho-IκBα, and β-tubulin. Inducers are 10 ng/ml TNFɑ (T), 10 µM camptothecin (C), 10 µM VP16 (V), 25 µM doxorubicin (D) and 5 Gγ irradiation (I). (B) Clone 1 cells were transfected with pcDNA-FLAG or pcDNA-FLAG-NEMO. Two days later, cells were treated with TNFα (+) or were left untreated (-). Western blotting of whole-cell extracts was done for phospho-IκBα or for NEMO (top) or nuclear extracts (bottom) were probed for NF-κB p65. ns, non-specific band. Ponceau staining of the filters was performed to ensure approximately equal loading of protein extracts.

To confirm that the lack of responsiveness of clone 1 cells to NF-κB-activating agents was due to the loss of NEMO expression, we transfected clone 1 cells with an expression vector for FLAG-NEMO (which lacks the 5’ exon 1B sequences and is therefore not susceptible to gRNA targeting), and then treated the cells with TNFα. As shown in Fig. 3B, re-expression of NEMO in clone 1 cells restored TNFα-induced phosphorylation of IκBα as well as nuclear translocation of NF-κB subunit p65. Taken together, these results indicate that clone 1 cells are specifically defective for stimulus-based activation of canonical NF-κB signaling, and that TNFα-induced activation of NF-κB signaling in clone 1 cells can be restored by re-expression of NEMO.

### Reduced genomic heterogeneity in a subclone of clone 1 cells

Because of the heterogeneity of CRISPR/Cas9-directed genomic deletions at the *NEMO* core promoter site in clone 1 cells, we sought to isolate a cell clone with a reduced number of genomic disruptions at the targeted site. Therefore, we made cell subclones by plating clone 1 cells at less than one cell per well. Five such cell subclones were then expanded and analyzed. Like parental clone 1 cells, all subclones expressed no detectable NEMO protein (Fig. 4A). Preliminary sequence analysis indicated that clone 1.1 had the least genomic complexity around the targeted site, and so clone 1.1 was the focus of further analysis. As shown in Fig. 4B, analysis of ~16,000 sequence reads of a PCR product covering the genomic disruption of clone 1.1 cells showed that over 99% of the reads contained an 8-bp deletion at the targeted site (and about 0.9% of the reads had a 1-bp deletion of an A residue in the triplet of the consensus sequence, i.e., ACCGCG-AACT). A second cell subclone (clone 1.3) had over 80% of its total reads (9,485) showing a 2-bp deletion in the AAA stretch of the consensus sequence (ACCGCG--ACT). As a second method of confirming the genomic editing at the target site in clone 1.1 cells, we used PCR amplification of the targeted site followed by restriction digestion with BsiEI, as there is a BsiEI restriction enzyme site at the wild-type target sequence (see Fig. S2) that would be destroyed by the 8-bp deletion in clone 1.1 genomic DNA. As predicted, the PCR product amplified from control 293T cell genomic DNA was digested by BsiEI, but the PCR product from clone 1.1 cell genomic DNA was not digested by BsiEI (Fig. 4C).

**FIG 4.**
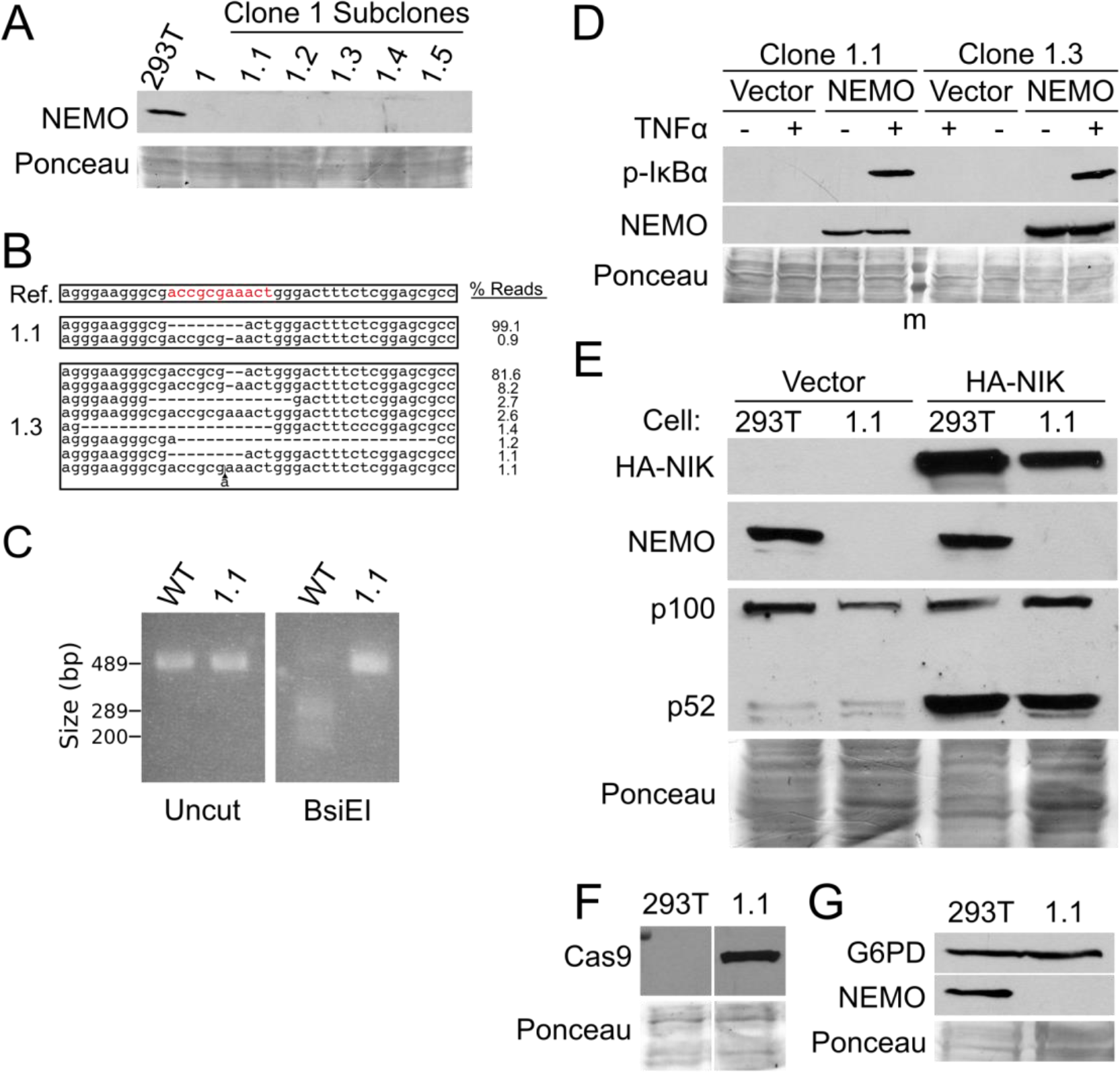
Isolation of cell subclones from clone 1 cells. (A) Five subclones of clone 1 cells were isolated and analyzed by Western blotting for NEMO. Shown also are control 293T and clone 1 cell extracts. (B) CRISPR sequencing of the PCR-amplified exon 1B core promoter in clones 1.1 and 1.3 was used to characterize the genomic disruptions (as done for Fig. 2C). (C) PCR-amplified exon 1B core promoter isolated from WT 293T and clone 1.1 cell genomic DNA was analyzed as Uncut or BsiE1-digested DNA. DNA was electrophoresed on a 1.5% agarose gel and was detected with ethidium bromide. (D) Clone 1.1 and 1.3 cells were transfected with pcDNA-FLAG or pcDNA-FLAG-NEMO. Two days later, cells were treated with TNFα (+) or were left untreated (-). Whole-cell extracts were analyzed for phospho-IκBα or NEMO expression. (E) Control 293T cells and clone 1.1 cells were transfected with pcDNA vector control or pcDNA-HA-NIK. Whole-cell extracts were subjected to Western blotting for HA-NIK, NEMO, and p100/p52. (F) Control 293T cells and clone 1.1 cell extracts were analyzed by Western blotting for FLAG-Cas9 (anti-FLAG). (G) Extracts from control 293T and clone 1.1 cells were analyzed by Western blotting for G6PD and NEMO. Where indicated, Ponceau staining of the filters was done to ensure approximately equal loading of total protein.

To characterize further the 1.1 and 1.3 cell subclones, we first assessed whether IκBα was phosphorylated in these cells in response to TNFα. As shown in Fig. 4D, 1.1 and 1.3 cells transfected with an empty vector did not show phosphorylation of IκBα following treatment with TNFα, whereas clone 1.1 and 1.3 cells transfected with a NEMO expression vector showed readily detectable TNFα-induced phosphorylation of IκBα.

Activation of non-canonical NF-κB signaling, which occurs by IKKα-directed phosphorylation and processing of NF-κB p100, has generally been found to be independent of NEMO (Sun, 2011). Consistent with these findings, overexpression of NIK, an activator of non-canonical NF-κB signaling, led to equal levels of NF-κB p100 processing to p52 in both wild-type 293T and clone 1.1 cells (Fig. 4E). Of note, the FLAG-Cas9 protein is still expressed in clone 1.1 cells, even after subcloning and extensive passaging (Fig. 4F).

### HDR-mediated repair of the 8-bp deletion in 1.1 cells

To demonstrate that NEMO lesions can be repaired, we attempted HDR-mediated repair of the 8-bp deletion locus in clone 1.1 cells. To do so, we co-transfected 1.1 cells with an expression vectors for a gRNA-directed against the site that was created by the 8-bp deletion and for a single-stranded repair oligo that also had a mutation (GGG->GAG) which abolished the adjacent PAM site used for the gRNA (see Table S2). The PAM site mutation was included so that the original gRNA and Cas9 in these cells could not target the repaired allele and so that we could confirm that the repair had occurred (and was not, for example, due to contamination). PCR amplification and BsiEI digestion of the targeted locus from HDR-transfected 1.1 cells showed that approximately 5-10% of the PCR-amplified product was digested with BsiEI (that is, a titration of wild-type DNA, indicated that approximately 5-10% of the PCR product from HDR-transfected 1.1 cells was digested with BsiEI [not shown]). Moreover, next generation sequencing showed that 11.5% of ~40,000 sequence reads had the wild-type sequence in place of the 8-bp deletion and also had the introduced GAG mutation in the adjacent PAM site (Fig. S3).

### Expression of *G6PD* from the bi-directional promoter is not affected in clone 1.1 cells

The *NEMO* exon 1B promoter is a bi-directional GC-rich (TATA box-lacking) promoter (Galgóczy et al., 2001; Fusco et al., 2012). That is, on the opposite DNA strand in the opposite direction, there are multiple transcriptional start sites for the *G6PD* gene. Western blotting showed that expression of glucose-6-phosphate 1-dehydrogenase protein is not altered in clone 1.1 cells as compared to control cells (Fig. 4G). Therefore, the 8-bp deletion in clone 1.1 cells is unlikely to have an effect on *G6PD* transcription, indicating that the 8-bp deletion inactivates the bi-directional promoter only for *NEMO* expression.

### Treatment of 293T cells with the gRNA against the exon 1B core promoter element and dCas9 also reduces NEMO expression

Based on the above experiments, we reasoned that other methods of targeting the exon 1B region might result in reduced NEMO expression. In particular, targeting of genomic regions with dCas9, which has mutations in its catalytic residues required for DNA-cleaving activity, has been shown to block specific gene expression under some conditions (Lawhorn et al., 2014; Fulco et al., 2016). Therefore, we first created a version of pLenti-CRISPR2.0 in which Cas9 was mutated at residues 10 and 840 (D10G and H840A), which inactivates the DNA cleaving function of Cas9 (Qi et al., 2013). As above, we then selected 293T cells that had been transduced with the pLenti-CRISPR2.0-dCas9 vector containing the *NEMO* exon 1B gRNA. We then measured NEMO expression by Western blotting in pools and clones of puromycin-selected cells. As shown in Fig. 5A, NEMO expression was reduced in two of three pLenti-CRISPR2.0-dCas9 cell clones containing the exon 1B gRNA and in a pool of transduced cells, as compared to the uninfected control 293T cells. As a control, no NEMO protein was detected in a lysate from clone 1.1 cells (Fig. 5A). The extent of NEMO protein knockdown in the pool of CRISPR2.0-dCas9 cells containing the NEMO exon 1B gRNA was similar to that seen in cells with the exon 1B gRNA and wild-type Cas9 (compare Fig. 5A to Fig. 2A). As a further control, we show that the dCas9 protein was expressed in cell clones d2 and d3. Moreover, no dCas9 protein was expressed in clone d1, likely explaining why there was no reduction in NEMO protein in those cells (Fig. 5A). In addition, G6PD protein expression was not affected in clones or pools of cells with reduced NEMO protein from dCas9/gRNA transduction (Fig. 5B). Finally, we sequenced the gRNA target site in the genomic DNA from clones d2 and d3. As expected, none of the amplified products for clones d2 and d3 (0 out of 47,208, and 23,507, respectively; data not shown) had genomic alterations at the target site, indicating that the dCas9 protein expressed in these clones is defective for genome editing. Taken together, the results in this section indicate that occupation of the *NEMO* exon 1B core promoter target site by the gRNA-dCas9 complex blocks efficient transcription of NEMO in 293T cells.

**FIG 5.**
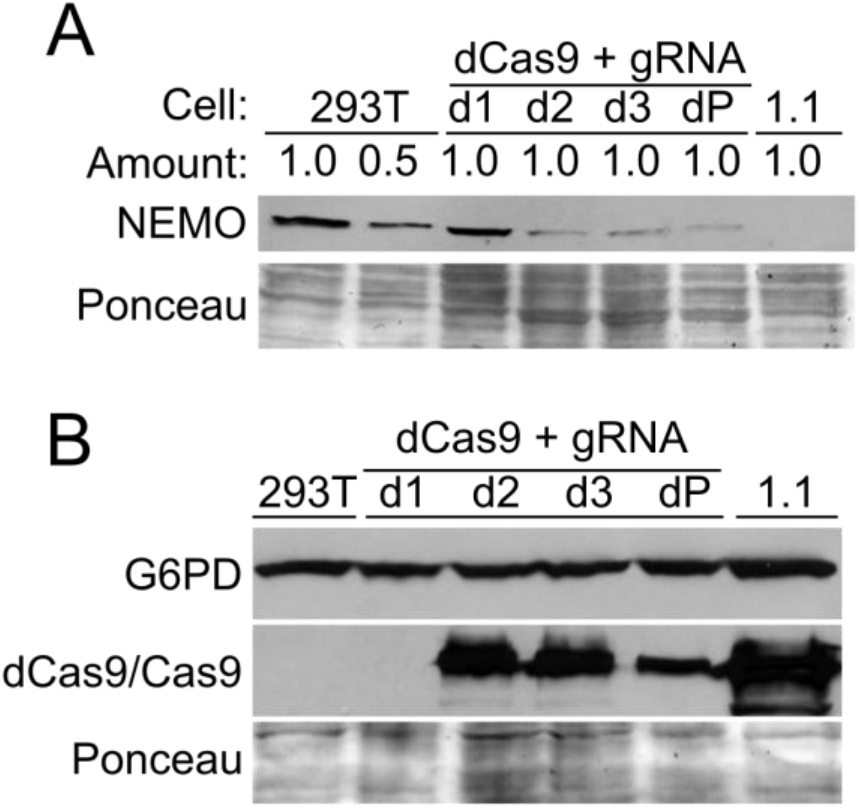
Targeting of the *NEMO* exon 1B core promoter with CRISPR/dCas9 reduced NEMO expression in 293T cells. (A) 293T cells were infected with a LentiCRISPR2.0-dCas9 construct containing exon 1B gRNA, and transduced cells were selected with puromycin. Western blotting for NEMO was then performed on control cells (positive control 293T and 50% of the amount of 293T cell lysate, or negative control clone 1.1 cells) or dCas9-infected pools (dP) or single cell clones (d1, d2, d3). (B) The indicated cell extracts were analyzed by Western blotting for G6PD and Cas9/dCas9. In both cases, Ponceau staining of the filters was done to ensure approximately equal loading of total protein.

### CRISPR/Cas9 targeting of the *NEMO* exon 1B core promoter element does not abolish NEMO expression in a human liver cell line

Fusco et al. (2006) reported that the exon 1D promoter is the major promoter used for *NEMO* transcription in liver cells, with approximately 14-fold higher expression of mRNA from exon 1D than exon 1B in both adult human liver tissue and the HepG2 liver cell line. Therefore, we hypothesized that CRISPR-Cas9-based targeting of the exon 1B core promoter sequence would not substantially affect NEMO expression in liver cells. To test this hypothesis, we transduced SNU-423 human liver cells with our pLenti-Crispr2.0 gRNA-containing vector and selected cells with puromycin. As shown in Fig. 6A, a puromycin-resistant clone of SNU-423 cells (clone L1) transduced with the gRNA vector expressed levels of NEMO protein that were essentially the same as the parental SNU-423 cells. As a control, we show that the clone LI cells expressed Cas9 (Fig. 6A). In addition, BseE1 digestion of the targeted site in the L1 clone (Fig. 6B) and DNA sequencing of the targeted locus in exon 1B showed that approximately 96% of the ~41,000 sequence reads were disrupted by genome editing (Fig. 6C). qPCR showed that clone L1 cells expressed approximately 60-80% of the levels *NEMO* mRNA as parental SNU-423 cells (Fig. 6D). Western blots of four additional Lenti-Crispr2.0 gRNA-containing SNU-423 cell clones showed that they expressed similar amounts of NEMO as compared to control cells (Fig. S4). Overall, these results indicate that targeting of the *NEMO* exon 1B promoter does not substantially affect usage of the upstream liver-specific *NEMO* promoter 1D, and thus, does not dramatically reduce NEMO protein expression in a human liver cell line.

**FIG 6.**
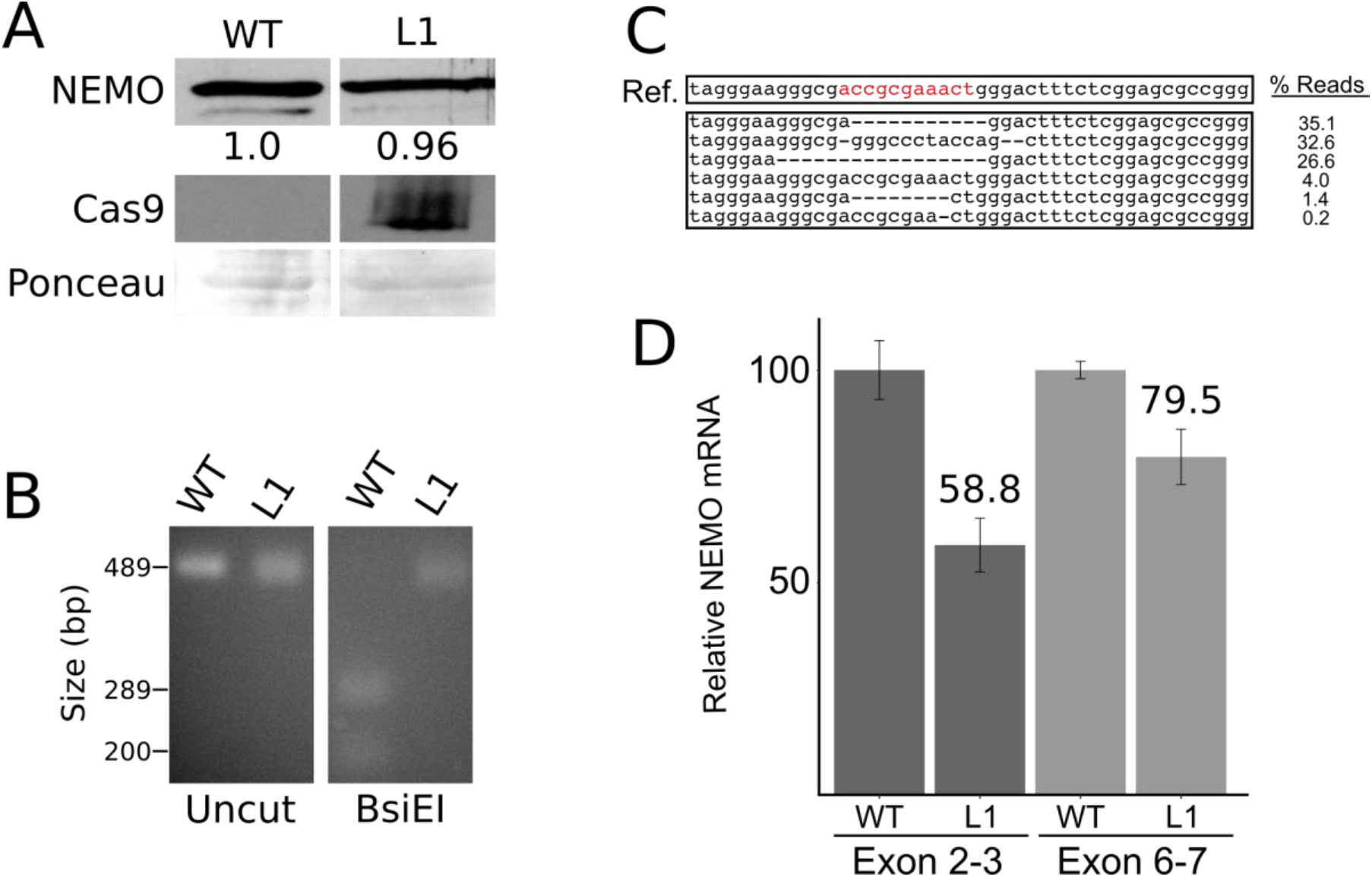
Targeting of the *NEMO* exon 1B core promoter with CRISPR/Cas9 does not affect NEMO expression in liver cells. (A) SNU-423 cells were transduced with a LentiCRISPR2.0-Cas9 construct containing exon 1B gRNA, and transduced cells were selected with puromycin. A cell clone was picked and expanded. Western blotting for NEMO and Cas9 was then performed on wild-type (WT) control cell or the liver cell clone (L1) extracts, or the filter was stained with Ponceau for total protein. The numbers below the NEMO lanes indicate the relative amount of NEMO protein in the WT and LI lanes, as determined by scanning of the film with ImageJ. (B) PCR-amplified exon 1B core promoter from WT SNU-423 and clone L1 genomic DNA was analyzed as Uncut or BsiE1-digested DNA. DNA was electrophoresed on a 1.5% agarose gel and detected with ethidium bromide. (C) CRISPR sequencing of the PCR-amplified exon 1B core promoter in L1 genomic DNA was used to characterize the genomic disruptions (as done for Fig. 2C.). (D) qPCR of *NEMO* transcripts was performed on cDNA isolated from WT and LI clone SNU-423 cells, using primers for exons 2 and 3 or 6 and 7 (as described for Fig.

## DISCUSSION

In this paper, we have shown that CRISPR-based targeting of a core promoter region of the most commonly used promoter of the *NEMO* gene can efficiently knockdown NEMO protein expression and function in human HEK 293T cells. Moreover, targeting of this core promoter sequence only marginally affects NEMO expression in a human liver cell line, in which *NEMO* transcription is controlled by a distinct liver-specific promoter. Although other researchers have targeted gene regulatory regions by CRISPR-based screening (e.g., Sanjana et al., 2016), to our knowledge, this is the first use of CRISPR-based genetic editing to ablate expression of an individual gene in a cell type-specific manner.

Transcription of the *NEMO* gene in the majority of cell types is controlled by a GC-rich, bi-directional promoter (promoter B) that has multiple transcriptional start sites (Fusco et al., 2012), as is commonly found with GC-rich promoters (Vo Ngoc et al. 2017b). The gRNA that we used to knockdown *NEMO* expression in 293T cells targets exon 1B core promoter sequences that are just downstream of a major *NEMO* TSS. Of note, this gRNA covers a sequence (ACCGCGAAACT) that is similar to one of four consensus sequences found near, and often downstream of, many eukaryotic/human TSSs (Vo Ngoc et al. 2017a) (Fig. 1A). This sequence is likely to be important for the initiation of transcription due to binding of a basal transcription protein or complex (Vo Ngoc et al., 2017a, 2017b). Consistent with the core sequence in the exon 1B promoter being a binding site for a transcription complex, we also found that targeting of dCas9 to this element blocks NEMO protein expression (Fig. 5). Although the consensus sequence logo for this 11-bp common promoter element is quite variable (Fig. 1A), it has three invariable A residues. Of note, all 42 of the genomic deletions in our initial 293T cell clone 1 have disruptions in this AAA sequence, including ~28% of the sequence reads that lack only a single A residue. Given that *NEMO* mRNA levels were reduced by at least 50-fold in clone 1 cells (Fig. 2D) and NEMO protein was essentially undetectable (Fig. 3A), it is unlikely that there is any substantial *NEMO* mRNA expression from the ~28% of *NEMO* alleles that have the 1-bp deletion in the clone 1 population. Moreover, approximately 90% of the edited alleles in clone 1.3 cells had 1 or 2 bp deletions in the AAA sequence (Fig. 4B) and expressed no detectable NEMO protein (Fig. 4A).

The clone 1 cells were established from a cell clone that was generated on the first transduction step, and yet clone 1 cells had a highly diverse set of at least 42 disruptions at the targeted site (Fig. S1). We believe that the target site heterogeneity within this “clone” (clone 1) arose from at least two sources: 1) that not all *NEMO* alleles were targeted in the originally transduced cell and therefore, additional disruptions occurred after the first cell division; and 2) that some early arising targeted alleles underwent further editing after the first targeting event (e.g., a 1-bp deletion could have been further edited to an 8-bp deletion), since these cells stably express the *NEMO* gRNA and Cas9. In contrast, the subclone 1.1 cells were isolated some time after passage of the original clone 1 cells, and thus the *NEMO* alleles (and the 8-bp deletion) in 1.1 cells were likely stabilized in these clonally re-derived cells. In any event, the clone 1.1 cells (derived from the clone 1 cells) represent a nearly homogenous edited cell line that will likely be useful for researchers who seek to investigate NEMO protein function.

One common method to analyze NEMO mutant function is to re-express the NEMO protein in mouse *NEMO* knockout cells (Schröfelbauer et al., 2012; Cote et al., 2013). Such experiments have two limitations: 1) human NEMO proteins are being analyzed in mouse cells, and 2) to establish pure, selected populations of NEMO reconstituted cells can take a month or more. The 1.1 cell line, which has a nearly homogenous and defined 8-bp deletion in exon 1B (Fig. 4B), is likely to be useful for the rapid analysis of NEMO mutants, as occur in human disease or created in the lab. That is, in an experiment that took less than a week, we showed that transient transfection of 1.1 cells with FLAG-NEMO can restore TNFα-induced phosphorylation of IκBα (Fig. 4D).

Many cancers rely on canonical NF-κB signaling for growth and survival (Paur et al, 2018). Relevant to our study, a CRISPR-based screen found *NEMO* to be among the top 7% of targets for restricting the growth of B-lymphoma cells (Reddy et al., 2017), and lymphoid cells use promoter B for expression of *NEMO* (Fusco et al., 2006). Almost all approaches that have sought to target NF-κB function have been directed at NF-κB pathway protein targets (Gilmore & Garbati, 2011). The limitations of such protein-based approaches is that, without an efficient organ-specific delivery system, one will affect the NF-κB in non-cancer cells. In particular, such NF-κB-direct approaches have often resulted in liver toxicity because targeting of NF-κB signaling, and even disruption of *NEMO* itself, causes liver toxicity (Luedde et al., 2007; Grivennikov et al., 2010; Luedde & Schwabe, 2011). Our finding that gRNA-directed targeting of *NEMO* promoter B does not substantially affect liver-specific (promoter A) expression of NEMO (Figs. 6 and S4) suggests that targeting of the core promoter B sequence for the treatment of cancers that rely on canonical NF-κB signaling would not affect liver function *in vivo*. Precisely how one would target *NEMO* promoter B *in vivo* with CRISPR/Cas9 is not obvious; however, several gRNA/Cas9 *in vivo* delivery systems have been investigated (Dunbar et al., 2018; Glass et al., 2018).

Finally, we have also shown that we can repair a *NEMO* mutation by HDR. That is, by HDR we provide evidence that we were able to correct the 8-bp promoter deletion in clone 1.1 cells (Fig. S3). This result establishes a precedent for repairing the small genomic alterations in *NEMO* that have been found in human patients (Maubach et al., 2017).

Overall, we believe the methods, results, and cell lines generated in this paper will be of interest to others studying NEMO function, and can provide novel methods of gene targeting (possibly for therapeutic purposes) and for investigating *in vivo* effects of alternative promoter usage.

## MATERIALS AND METHODS

### Cell culture and plasmids

Human embryonic kidney 293T (HEK 293T) and SNU-423 cells (a gift of Ulla Hansen, Boston University) were cultured in Dulbecco’s Modified Eagle Medium (DMEM; Thermo Fisher) with 10% Fetal Bovine Serum (FBS; Biologos), supplemented with 50 U/ml penicillin, and 50 µg/ml streptomycin) at 37 °C, 5% CO2, in a humidified incubator.

Detailed descriptions of all plasmids and primers used in this paper are presented in Tables S1 and S2. Plasmids pcDNA-FLAG, pcDNA-FLAG-NEMO have been described previously (Herscovitch et al., 2008). pLentiCRISPRv2.0 is from Addgene (plasmid # 52961; Sanjana et al., 2014). The gRNA sequences targeting the *NEMO* exon 1B core promoter (quality score = 96; 39 off-target sites; crispr.mit.edu) or a control sequence (labelled C in Fig. 2B) slightly downstream of the exon 1B site (Table S2) were synthesized with overhangs and were ligated into BsmBI-digested plentiCRISPRv2.0. pLentiCRISPRv2.0-dCas9 plasmid was constructed using PCR site-directed overlap mutagenesis in two steps: first, both D10G, and H840A mutations were introduced via amplification with overlapping primers spanning the N, and C-terminal halves of dCas9, the resulting two fragments containing the N- and C-terminal halves of dCas9 were subcloned stepwise into a pSL1180 carrier plasmid. Finally, full-length dCas9 was then subcloned into the LentiCRISPR v2.0 plasmid, and confirmed by DNA sequencing. The relevant portions of plentiCRISPRv2.0-gRNA and pLentiCRISPRv2-dCas9 plasmids containing inserts were also verified by DNA sequencing.

### Establishment of CRISPR *NEMO* targeted cell lines

293T cells were seeded at ~60% confluence in a 100-mm tissue culture plates approximately 24 h prior to transfection. The next day, cells were co-transfected with 1 µg plentiCRISPR v2, 2 µg pCMV-dR8.91, and 4 µg pCMV-VSV-G along with 32 µg polyethylenimine (PEI) in a 1 ml serum-free DMEM, and incubated for 15 min at room temperature. The PEI:DNA-containing transfection mixture was diluted to 10 ml with DMEM/10% FBS, added to plates of 293T cells, and incubated overnight for approximately 16 h. On the next day, plates were washed once with PBS, and fluid changed with fresh DMEM/10% FBS. Cells were then incubated for two more days, and at that time, transfected cells were subcultured into new 100-mm plates with 10 ml DMEM/10% FBS. The following day, puromycin was added to a final concentration of 2.5 µg/ml. Plates were maintained under puromycin selection, and fluid changed approximately every 3-5 days, until distinct colonies had formed (at approximately 15 days after start of puromycin selection). Cell lines were established from single puromycin-resistant, and were screened for NEMO expression by Western Blotting (e.g., clone 1). After approximately eight passages, subclones of clone 1 cells were isolated by limiting dilution. That is, clone 1 cells were plated in a 96-well tissue culture plate at a density of 0.25 cells/well, and clones (e.g., 1.1 – 1.5) growing out of individual wells were re-screened for NEMO expression by Western blotting.

### Western blotting

Whole-cell extracts were prepared in AT lysis buffer (20 mM HEPES pH7.9, 150 mM NaCl, 1mM EDTA, 1 mM EGTA, 20% w/v glycerol, 1% v/v Triton X-100, and Roche Complete Protease Inhibitor). Extracts were then separated on a 10% SDS-polyacrylamide gel, transferred to a nitrocellulose membrane in transfer buffer (20 mM Tris, 150 mM glycine, 20% methanol) overnight at 4 °C. To detect high molecular weight Cas9 and dCas9 proteins, whole-cell extracts were resolved on a 7.5% SDS-polyacrylamide gel, and transferred to nitrocellulose in low-methanol transfer buffer (20 mM Tris, 150 mM glycine, 20% methanol) at 250 mA for 1 h, followed by 170 mA overnight under 4 °C. Membranes were blocked in TMT (10 mM Tris-HCl pH 7.4, 150 mM NaCl, 0.1% v/v Tween 20, 5% w/v Carnation nonfat dry milk) for 1 h at room temperature, incubated in the appropriate primary antibody overnight at 4 °C as follows: anti-NEMO (1:1000; #2689, Cell Signaling Technology), anti-phospho-IκBα (1:1000; #9246, Cell Signaling Technology), anti-FLAG (1:1000; #2368, Cell Signaling Technology), anti-HA (1:500; Y-11, Santa Cruz Biotechnology), anti-NF-κB p100 (1:500; #4882, Cell Signaling Technology), anti-p65 (1:2000, #1226 gift of Nancy Rice), anti-tubulin (1:500), anti-G6PD (1:500, #12263, Cell Signaling Technology), and anti-Cas9-HRP (1:10,000; ab202580, Abcam). Membranes were washed four times with TBST (10 mM Tris-HCl pH 7.4, 150 mM NaCl, 0.1% v/v Tween 20). Membranes were then incubated with the appropriate secondary HRP-linked antibody as follows: anti-rabbit-HRP (1:3000, Cell Signaling Technology) for NEMO, FLAG, Tubulin, p65, and G6PD or anti-mouse-HRP (1:3000, Cell Signaling Technology) for anti-phospho-IκBα. Membranes were then washed three times with TBST, twice with TBS (10 mM Tris-HCl pH 7.4, 150 mM NaCl), and incubated for 5 min at room temperature with chemiluminesent HRP substrate (SuperSignal West Dura Extended Duration Substrate, Thermo Fisher). Immunoreactive bands were detected by exposing membranes to autoradiography film (GeneMate Blue Basic Autoradiography Film, BioExpress).

### Genomic DNA amplification and analysis

Genomic DNA was isolated from cultured cells by first lysing the cells in a buffer consisting of 10 mM Tris-HCl, pH 8, 100 mM NaCl, 25 mM EDTA, 0.5% SDS, 1 mg/ml proteinase K. Samples were then heated for 2 h at 60°C, extracted with phenol two times and with chloroform once. DNA was precipitated with 100% ethanol for 30 min. Approximately 0.5 µg of genomic DNA was used in PCRs. For DNA sequence analysis of the targeted site, DNA was amplified by standard PCR using primers NEMO gExon 1B Fwd and NEMO gIntron 1 Rev (see Table S2). The PCR product was purified and then sent out for Next-Generation sequencing (MGH CCIB DNA Core). For restriction enzyme analysis of the edited site, DNA was amplified by standard PCR using primers gDNA Fwd and gDNA Rev (Table S2). From a 50 µl reaction, 7 µl was then digested with BsiEI (New England Biolabs) and analyzed by agarose gel electrophoresis followed by staining with ethidium bromide.

### HDR-mediated repair of the mutated NEMO allele in clone 1.1 cells

Clone 1.1 cells were seeded at ~60% confluence in 35-mm tissue culture plates approximately 24 h prior to transfection. The next day, cells were co-transfected with 1.8 µg of px330puro-HDR (see Table S1), 0.2 µg of an 84-bp single-stranded HDR template (HDR-1.1; Table S2), along with Effectine reagents (QIAGEN; see below) in DMEM containing 10% FBS. On the next day, the transfection media was replaced with fresh DMEM/10% FBS. Two days later, cells were passaged into 60-mm plates with 5 ml DMEM/10%FBS. After allowing the cells to grow out, genomic DNA was isolated and target DNA was amplified by standard PCR using primers NEMO gExon 1B Fwd and NEMO gIntron 1 Rev (see Table S2). The PCR product was purified and subjected to Next-Generation sequencing (MGH CCIB DNA Core).

### RNA analysis by RT-PCR and qPCR

mRNA was isolated from cultured cells using TRIzol RNA purification kit (QIAGEN) according to the manufacturer’s instructions. cDNA was synthesized using random primers and M-MLV reverse transcriptase or the TaqMan cDNA synthesis kit (Thermo Fisher). For RT-PCR, *NEMO* cDNA was amplified by PCR for 32 cycles (using primers NEMO Exon 1B Fwd NEMO Intron 1 Rev [Table S2]), and DNA was then analyzed by polyacrylamide gel electrophoresis and staining with ethidium bromide. For qPCR, *NEMO* cDNA was amplified (using primers NEMO Exon 2 Fwd and NEMO Exon 3 Rev or NEMO Exon 6 Fwd and NEMO Exon 7 Rev) using SYBR Green Real-Time PCR kit (Thermo Fisher), and analyzed on an ABI 7900ht qPCR machine. The delta delta CT method was used to quantify the relative fold change in gene expression, and values were normalized to *NEMO* mRNA expression in control 293T or SNU-423 cells. For RT-PCR and qPCR *GAPDH* mRNA was used as a control (see primers in Table S2).

### Cell treatments

293T and clone 1 cells were treated with various NF-κB inducers (10 ng/ml TNFα for 30 min; 10 µM camptothecin for 2 h; 10 µM VP16 for 2 h; 25 µM doxorubicin for 2 h; 5 Gy of ionizing irradiation and terminated after 1.5 h. A JL Sheperd model JL-10 with a cesium (137CS) source was used for gamma-irradiation. Whole-cell extracts were made using total extract (TOTEX) buffer (20 mM HEPES (pH 7.9), 350 mM NaCl, 1 mM MgCl2, 0.5 mM EDTA, 0.1 mM EGTA, 0.5 mM DTT, 20% glycerol, and 1% NP-40). EMSAs and immunoblots were performed as described previously (Miyamoto et al., 1998; Jackson et al., 2015) and below. In Fig. 3A, the following antisera were used: anti-NEMO (FL-419; sc-8330 antibody; Santa Cruz); anti-tubulin (CP06; EMD Millipore); anti-phospho-IκBα (5A5; #9246, Cell Signaling)

### NEMO reconstitution experiments

Exon 1B-edited cells (clone 1, 1.1 or 1.3) were seeded at ~60% density in a 60-mm tissue culture plates one day prior to transecting with the pcDNA-FLAG empty vector or pcDNA-FLAG-NEMO. Transfections were performed using Effectene Transfection Kit (QIAGEN) as follows: 0.2 µg plasmid, and 1.6 µl Enhancer were diluted to a final volume of 32 µl with EC Buffer, and incubated for 5 min at room temperature for DNA condensation. Then, 2 µl Effectene was added, and samples were incubated for 7 min at room temperature to form Effectene-DNA complexes. The final transfection mixture was brought to 1.8 ml with DMEM/10% FBS, added to 60-mm plates containing 3.2 ml of fresh medium, and incubated overnight. Upon reaching confluence (2 days), transfected plates were subcultured into new plates at ~60% density, and incubated again until cells reached confluence (2 days). Cells were then stimulated with 20 ng/ml recombinant TNFα (R&D Systems) diluted in PBS containing 0.1% BSA for 10 min in a 37 °C, 5% CO2 in a humidified incubator. Cells were then lysed on ice and processed for Western blotting as described above.

### Nuclear extract preparation

To prepare nuclear fractions, cells were resuspended in 400 µl hypotonic buffer (10 mM HEPES pH 7.9, 1.5 mM MgCl2, 10 mM KCl), and were incubated on ice for 10 min. Samples were then supplemented with 55 µl NP-40, vortexed for 10 sec, and pelleted at 500 x g for 5 min at 4 °C. The nuclear pellet was washed with the hypotonic buffer, and re-pelleted as above. To lyse the nuclei, the pellet was resuspended in 60 µl high-salt buffer (20 mM HEPES pH 7.9, 1.5 mM MgCl2, 0.2 mM EDTA, 420 mM NaCl, 25% v/v glycerol), vortexed for 30 sec, and samples were then rocked for 1 h at 4 °C. The nuclear extract was clarified by pelleting debris for 30 min at 13,000 rpm at 4 °C.

## SUPPLEMENTAL MATERIAL

Two Supplementary Tables and four Supplementary Figures.

## ACKNOWLEDGMENTS

We thank Juan Fuxman Bass and Trevor Siggers for comments on the manuscript, and Demetrios Kalaitzidis for helpful discussions.

This research was supported by the National Institutes of Health grants CA077474 (to S.M.) and GM117350 (to T.D.G.). M.B., Y.L., L.K. and E.Z.M. were supported by funds from the Boston University Undergraduate Research Opportunities Program.

We have no conflicts of interest to declare.

